# Antibiotics in microbial communities: an ecological frame of resistance

**DOI:** 10.1101/2020.05.10.086744

**Authors:** Andrew D. Letten, Alex Hall, Jonathan Levine

## Abstract

There is growing awareness that our ability to tackle antibiotic resistance is limited by a lack of mechanistic understanding of the communities in which resistant microbes are embedded. The widespread coexistence of resistant and sensitive bacteria in microbial systems presents an especially frustrating paradox. Recent advances in ecological coexistence theory offer a powerful framework to probe the mechanisms regulating intra- and inter-specific coexistence, but the significance of this body of theory to the problem of antimicrobial resistance has been largely overlooked. In this Perspectives article, we draw on emerging ecological theory to illustrate how changes in both competitive ability and niche overlap are critical for understanding costs of resistance and the persistence of pathogens in microbial systems. We then show how trade-offs in resource acquisition strategies can have counter-intuitive consequences for the coexistence of resistant and susceptible genotypes in variable environments. These insights highlight numerous opportunities for innovative experimental and theoretical research into antibiotic resistance in the microbiome.

## Introduction

Less than a century since the discovery of antibiotics marked one of medicine’s greatest forward leaps, the evolution of resistance has emerged as one of its most pressing challenges^1^. To date, efforts to address this global health problem have primarily concentrated on restricting unnecessary prescriptions, the development of new drugs, and the use of novel combinations of existing drugs^2^. Now, however, with emerging insight into the critical role the host and environmental microbiomes play in regulating health and diseases, there is growing consensus that we need to look beyond one-to-one host-pathogen relationships and consider the complex web of competitive interactions within which individual pathogens are embedded^3–8^. This expanded ecological perspective holds promise not only for our fundamental understanding of resistance evolution, but also for the identification and development of novel treatment strategies.

A core challenge in tackling antibiotic resistance is understanding the balance between the selective advantage conferred by a resistance mutation and the selective disadvantage incurred by any associated fitness trade-offs, i.e. so-called costs of resistance^9,10^. Simple population models predict that because resistant and sensitive strains compete within the same habitat (e.g. within hosts or the environment), the fitter strain – determined by the level of antimicrobial exposure and the costs of resistance – should exclude the weaker strain ^11^. In reality, however, an unexpected observation is that sensitive and resistant strains commonly coexist even though they compete within hosts^11–13^, and that resistant strains often persist long after antibiotic exposure has ceased^14^. Not knowing how and why resistant and sensitive strains commonly coexist is widely recognised to be a major barrier to predicting the prevalence of resistance^11,15,16^, and ultimately prolonging the effective therapeutic life of antimicrobials^14^.

By convention, costs of resistance are defined as a decrease in the fitness (e.g. per-capita growth rate or yield) of a mutant strain possessing a resistant allele relative to an isogenic, ancestral strain in an antibiotic free environment^9,10^. In practice, costs of resistance are usually evaluated based on comparisons of demographic parameters in monoculture, or preferably based on the relative frequency of the mutant strain and the ancestral strain after 24-hours competing in batch culture^17^. Thus, ecologically speaking, a cost of resistance is analogous to a loss in competitive ability. Competitive ability is of course highly context dependent; a good competitor in one environment may be a poor competitor in another. Indeed, a number of studies have demonstrated the environmental contingencies of resistance costs^18–22^. Perhaps less obvious, and certainly much less studied, is the extent to which competitive ability depends on biotic context beyond the focal bacteria, i.e. the identity of community members and the propensity for competitive interactions between them. A large loss in competitive ability may be negligible in a depauperate community free from competitors, while conversely a small loss in competitive ability may have significant implications for survival in a more healthy, diverse system.

In the ecological literature, a rich body of theory has built up over the last 30-40 years with the primary goal of understanding the dependence of pairwise competitive outcomes on the broader community of interacting species^23–31^. Through a quantitative partitioning of differences in competitive ability and niche overlap between competing species, coexistence theory (as it is popularly termed) has been especially instrumental in crystallizing the fundamental rules of diversity maintenance and providing a mathematical framework to investigate the relative importance of deterministic and stochastic processes for coexistence in empirical systems^24,29,30^. In addition, it has been invaluable in disentangling contradictory perspectives on competitive ability and coexistence in equilibrium and non-equilibrium systems ^32–34^. While this framework has been embraced by community ecologists focusing on interspecific interactions, when reproduction is primarily clonal, as it is in many bacteria, the framework can be applied equally robustly to intraspecific interactions (i.e. competition between different genotypes within a species).

Despite the importance of ecological theory for understanding the evolution of resistance, the separation of these two research topics into largely independent literature has slowed their synthesis. Several research groups have begun to bridge this gap, with a number of studies illustrating the significant role within-host ecology can play in regulating the evolution of resistance in complex communities^7,8,16,35^. Nevertheless, advances in our understanding of coexistence at the frontier of the field remain untapped. Our primary goal in this paper is therefore to highlight core concepts from coexistence theory that have the potential to inform our understanding of how antibiotic resistant strains are able to persist alongside susceptible strains and/or the broader microbial community. We begin by outlining the fundamental tenets of coexistence theory and the insights that can be gained from partitioning costs of resistance into effects on competitive ability and effects on niche overlap, before exploring the existing empirical rationale for the partition. In the next section, we look deeper into the coexistence of susceptible and resistant strains in constant versus variable environments, drawing on emerging ecological theory on the effects of equilibrium vs. non-equilibrium dynamics on competitive ability. In the penultimate section, we identify some of the key limitations of existing ecological theory as applied to antibiotic resistance evolution, and how theory might be developed to address these limitations. We finish by identifying what we perceive to be the most important theoretical and empirical gaps in our understanding of the community ecology of costs of resistance.

Note, this contribution is not intended as a fully comprehensive review of the potential application of ecological theory to the evolution of antibiotic resistance. We are also acutely aware that there are myriad other factors related to pathogen dynamics, host immunity and pharmacology that may overwhelm any of the patterns we predict based on comparatively simple models. Nevertheless, we hope that the ideas presented herein will provide a valuable new perspective to researchers working in the field of antibiotic resistance with limited exposure to recent advances in ecological theory.

### Partitioning costs of resistance: competitive ability and the niche

While coexistence theory embodies a broad body of theory aimed at characterising the diverse mechanisms regulating coexistence in spatio-temporally homogenous and heterogeneous environments, one of its defining features is to separate the contributions differences in competitive ability^†^ and niche overlap make to the relative fitness of a focal species and its competitors^24,26,29^. Competitive ability differences capture how well adapted two species are to their shared environment, whereas niche differences capture how much they overlap in their usage of their shared environment. It follows that coexistence between competitors requires that their differences in competitive ability do not exceed the stabilizing effect of their niche differences. At the extreme, if two species completely overlap in resource usage (and share the same predators) in space and time, then they would need to have identical competitive ability in order to coexist, and even then only neutrally. However, the more species differentiate in resource usage the greater the range of differences in competitive ability that are compatible with coexistence, until the other extreme is reached, where two species that have zero niche overlap (i.e. do not use any of the same resources in space and time) can coexist in spite of infinitely large differences in competitive ability (see Box 1).

#### Box 1

Separating competitive ability and the niche

Although differences in competitive ability and niche overlap can hypothetically be obtained for any pairwise model of competition, the most convenient formulas, and the most readily used by empirical ecologists, are those derived for phenomenological Lotka-Volterra type competition models of the general form:

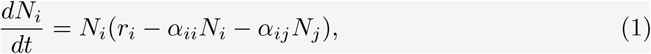

where *r*_*i*_ is the per capita intrinsic rate of increase of the focal species, *α*_*ii*_ is the linear effect of intraspecific competition (sometimes parameterized as carrying capacity, 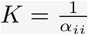), and *α*_*ij*_ is the linear effect of interspecific competition.

Niche overlap, *ρ*, is given by the geometric mean ratio of the intraspecific and interspecific coefficients,

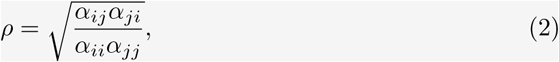

and the competitive ability ratio, 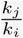, is given by the ratio of intrinsic growth rates multiplied by the geometric mean ratio of each species sensitivity to competition,

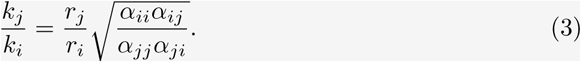

Based on the mutual invasibility criterion for coexistence, i.e., that each species can invade from rare when its competitor(s) is resident at its equilibrium density, stable coexistence requires that,

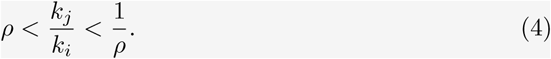

In other words, as niche overlap increases (approaches 1), an increasingly narrow range of differences in competitive ability are compatible with coexistence (see Figure 1b).

**Figure 1:**
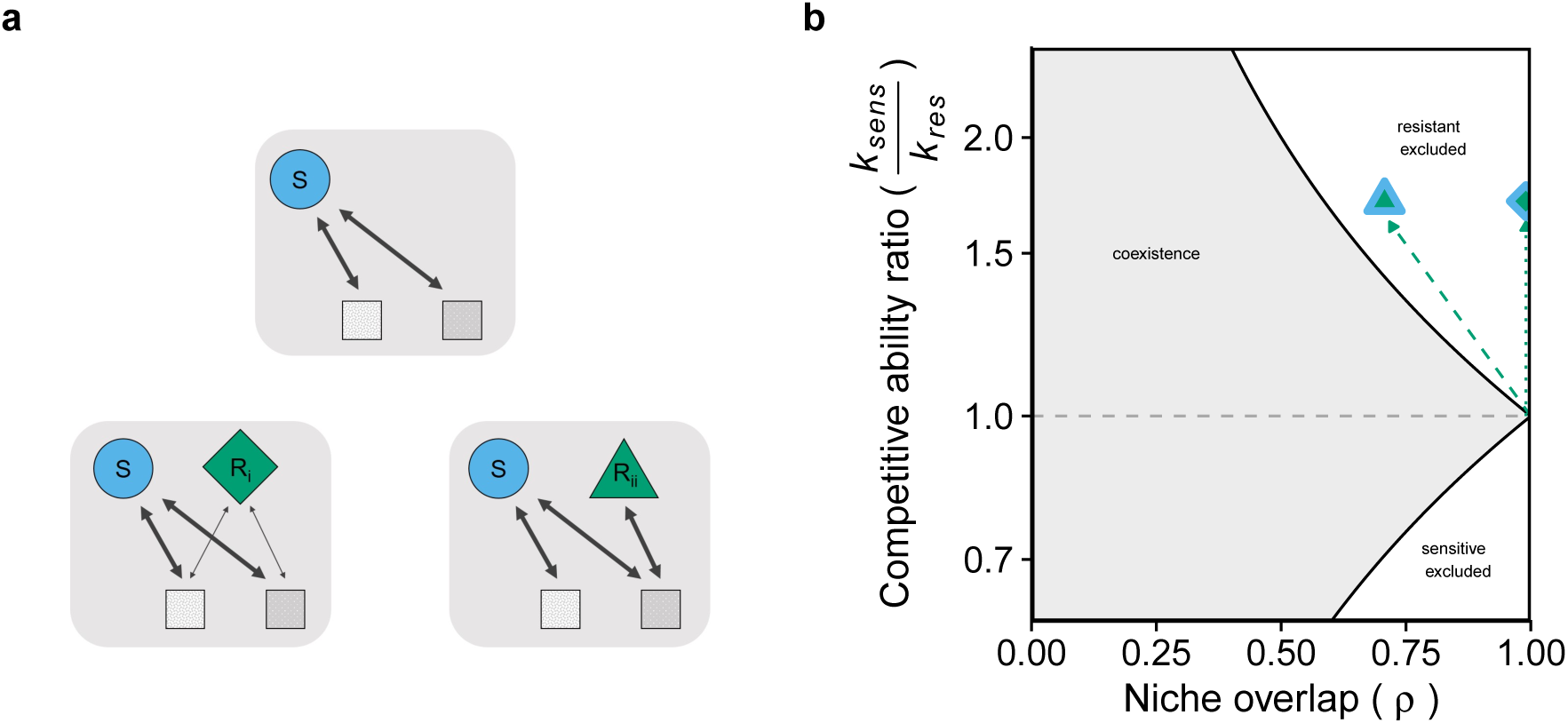
Partitioning costs of resistance mediated by constraints on resource uptake into competitive ability differences and niche overlap. **(a)** Basic food webs for a drug susceptible pathogen (*S*) consuming two substitutable resources in the absence (top) and presence (bottom) of two different drug resistant mutants, exhibiting either an equal magnitude loss in performance on both resources (*R*_*i*_, bottom left) or a loss in the ability to metabolize one of two resources with no change in performance on the other resource (*R*_*ii*_, bottom right). Line weight indicates consumer per-capita growth rate for each resource, and bidirectional arrows indicate that consumers and resources have reciprocal feedbacks on their respective densities. **(b)** Dotted and dashed green arrows trace the corresponding change in niche overlap and competitive ability differences between the susceptible and resistant strain (*R*_*i*_, green-filled blue diamond; *R*_*ii*_, green-filled blue triangle). Grey shaded region indicates parameter space corresponding to coexistence; unshaded region indicates parameter space corresponding to exclusion. See Supplementary Information S2 for model parameters.

For more mechanistic models of competition (e.g. consumer-resource models), which we use here to illustrate how costs of resistance can be partitioned into effects on competitive ability and niche differences, under certain assumptions we can obtain these metrics by obtaining mechanistic definitions for the Lotka Volterra parameters (see Supplementary Information S1). Competitive ability and niche differences then become explicit functions of the consumer growth and resource supply parameters, rather than the competition coefficients of Lotka-Volterra.

How then might separating relative fitness into competitive ability and niche overlap components inform our understanding of antibiotic resistance evolution? The critical insight is that costs of resistance may arise either solely from a loss in competitive ability or in conjunction with a change in niche overlap. Traditionally, costs of resistance have been interpreted primarily through the lens of a loss in competitive ability (i.e. a reduction in growth rate or yield on a given resource) ^17^. This might be reasonable, under certain assumptions, if we are only concerned with the relative fitness of a resistant mutant and its drug susceptible ancestor, but in the presence of the community the limitations of ignoring niche overlap come into especially sharp focus.

Consider a mutant that evolves resistance to an antibiotic resulting in either: reduced growth on all resources consumed by the ancestor, i.e. a general metabolic burden (*R*_*i*_, bottom left in Figure 1a); or zero growth on just a subset of resources consumed by the ancestor, i.e. more specific changes in bacterial physiology (*R*_*ii*_, bottom right in Figure 1a). The implications these different scenarios have on partitioning costs of resistance are illustrated in Figures 1&2 via the analysis of a classic mechanistic model of competition for two substitutable resources (see Supplementary Information S1 and S2 for details).

**Figure 2:**
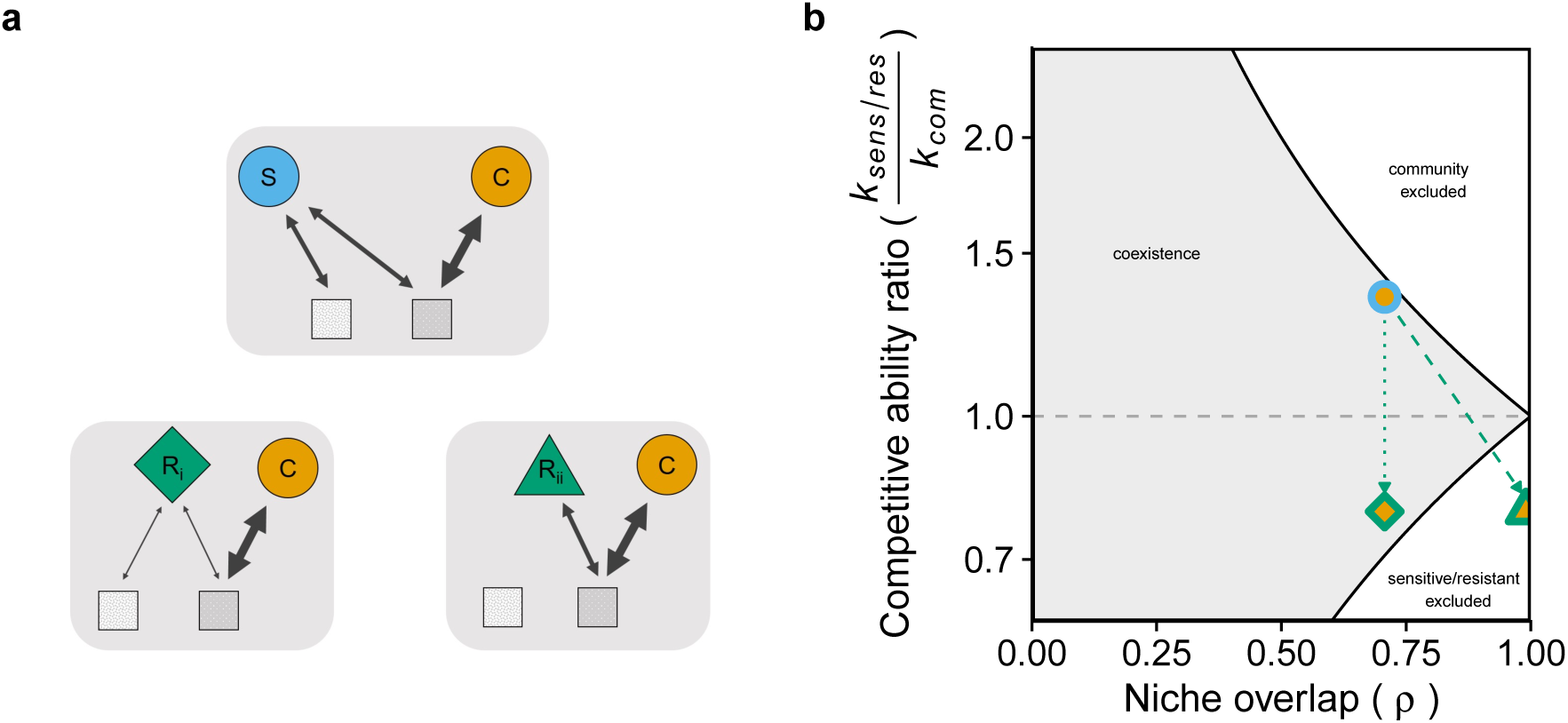
Partitioning costs of resistance mediated by constraints on resource uptake into competitive ability differences and niche overlap. **(a)** Basic food webs for a non-pathogenic member of the background community (*C*) consuming one of two substitutable resources in the presence of either an ancestral drug susceptible strain (top), or two alternative drug resistant mutants (bottom) exhibiting either an equal magnitude loss in performance on both resources (*R*_*i*_, bottom left) or a loss in the ability to metabolize one of two resources with no change in performance on the other resource (*R*_*ii*_, bottom right). Line weight indicates consumer per-capita growth rate for each resource, and bidirectional arrows indicate that consumers and resources have reciprocal feedbacks on their respective densities. **(b)** Orange-filled blue circle denotes the initial niche overlap and competitive ability differences between the susceptible strain and the background community member. Dotted and dashed green arrows trace the corresponding change in niche and competitive ability differences between the resistant strain and the background community member (*R*_*i*_, orange-filled green diamond; *R*_*ii*_, orange-filled green triangle). See Supplementary Information S2 for model parameters.

Despite very different trade-offs in their resource utilization traits, both mutant phenotypes exhibit identical costs of resistance in terms of reductions in their Malthusian growth parameters (growth rate at low density) and carrying capacities relative to the ancestral strain (see Supplementary Information S2). This is to say that the apparent costs of resistance of the two mutant phenotypes will be indistinguishable based on measurements of their growth in monoculture. From the perspective of coexistence theory, however, these costs of resistance present very different targets for control by commensal members of the background community.

Consider first the relationship between the ancestral strain and the two resistant phenotypes (Fig 1). Relative to the ancestral strain, a reduced ability to consume all resources (bottom left in Fig 1a and dotted green line in Fig 1b) will result in a reduction in competitive ability but no change in niche overlap. In contrast, losing the capacity to metabolize a previously consumed resource (bottom right in Fig 1a and dashed green line in Fig 1b) will not only result in the same reduction in competitive ability but also a reduction in niche overlap relative to the ancestral strain. In both cases, however, the outcome will be exclusion of the mutant strain by the susceptible in the drug free environment. It is nevertheless feasible that the mutant strain could gain access to (or better utilize) a resource that the ancestral strain does not utilize (or utilizes poorly), in which case the resultant change in niche overlap could be sufficient to foster the instantaneous coexistence of the resistant and susceptible phenotypes (see *Empirical basis*).

Now consider the relationship between the mutant phenotypes and a member of the background community. The significance of an increase in niche overlap between a drug resistant mutant and the background community depends to some extent on environmental context. In the absence of antibiotics, an increase in niche overlap between the drug resistant mutant and the background community could bring them into more direct competition, and therefore exaggerate observed costs of resistance (Fig 2b). This could be particularly important in preventing the evolution of resistance if the loss of competitive ability between the mutant strain and the ancestral strain is comparatively small. In contrast, in the presence of an antibiotic, any change in niche overlap between the mutant strain and the background community will be of little consequence if the members of the background community are also inhibited by the antibiotic. However, if some background community members are naturally insensitive to the antibiotic (e.g., in the case of narrow spectrum drugs that only target gram negative or positive bacteria), they could play a critical role in preventing a drug resistant mutant from dominating the system^8,16,35^.

Similarly, compensatory mutations that reduce the apparent costs of resistance can be usefully partitioned into their effects on competitive ability and their effects on niche overlap. For example, a compensatory mutation that reverses a resistance-associated decline in resource efficiency (i.e. a general metabolic burden), should equalize the competitive ability of the resistant and susceptible strain, and bring them closer to a state of potential neutral coexistence (a reversal of the dotted arrow in Figure 1b). The same would be true if a compensatory mutation restored the ancestral ability to metabolize a particular resource (a reversal of the dashed arrow in 1b). It is also possible, however, for a compensatory mutation to more strongly stabilize coexistence with its drug sensitive ancestor in spite of no change in the competitive ability of the resistant strain. This would arise if the resistant strain was to acquire access to a resource that the ancestral susceptible strain is unable to utilize.

#### Empirical basis

To our knowledge, no studies to date have looked directly at partitioning apart changes in competitive ability and niche overlap arising from resistance mutations. Nevertheless, there is significant evidence that resistance mutations are frequently associated with specific metabolic shifts rather than a more general metabolic burden^20,36–38^. For example, Perkins and Nicholson ^37^ tested substrate utilization patterns of strains of *Bacillus subtilis* with mutations in the RNA polymerase *β* subunit gene *rpoB* conferring resistance to the broad-spectrum antibiotic rifampicin. Given that single amino acid substitutions on RNA polymerase can influence expression across the entire genome^10^, it is perhaps not unexpected that they found that the uptake of several carbon substrates utilized strongly by the drug susceptible wild type was no longer detectable in many rifampicin resistant strains. This would be consistent with both a change in the competitive ability ratio and niche overlap between the wild type and resistant mutants as illustrated by the green-filled blue triangle in Fig 1b. More surprisingly, the authors also found that many mutant strains increased their utilization patterns on substrates utilized only weakly by the wild-type ^37^. The implication is that a single resistance mutation could actually result in a sufficiently large decrease in niche overlap to allow the susceptible ancestor and the resistant mutant to coexist, i.e., the green-filled blue triangle in Fig 1b would fall within the shaded coexistence region. Similarly, Paulander *et al.*^20^ *found that streptomycin resistant Salmonella typhimurium*, possessing a mutation on the ribosomal protein S12, were able to better utilize several (poorer) carbon sources than the susceptible wild type. Despite apparent fitness costs arising from reduced growth on favourable carbon sources, as well as an impaired stress response, this shift in resource utilization patterns may again be sufficient to allow streptomycin resistant mutants to coexist alongside the susceptible strain (assuming the availability of multiple carbon sources in their shared environment).

The previous examples suggest that chromosomal mutations that have pleiotropic effects on multiple traits are likely to result in both changes in competitive ability and niche overlap between wild type and resistant strains. Are there scenarios under which we might expect niche overlap to remain unchanged in spite of changes in competitive ability? One pathway by which fitness costs may exclusively affect competitive ability is when resistance is conferred through the overexpression of multidrug efflux pumps that result in resources being ejected before they can be used^39,40^. That being said, the overexpression of a multidrug efflux pump in *Stenotrophomonas maltophilia* has been shown to make resistant mutants more efficient in acquiring certain sugars and amino acids than the wild type ^36^. An alternative route to competitive-ability-centric fitness costs is plasmid carriage. The main fitness costs of plasmid carriage are thought to stem from the translation of protein-encoding plasmid genes, which diverts limiting resources and energy, including ATP, from other critical processes within the cell. This increased metabolic burden has been associated with reduced growth rates and cell densities, and lengthening of lag phases^41,42^. To the extent that these increased energetic costs will have broad cell-wide effects on metabolic fluxes and cellular processes ^41^, it may be expected that plasmid carriage will have minimal impacts on niche overlap despite significant fitness costs. Nevertheless, there are examples where plasmid carriage has been associated with very specific phenotypic changes, including the increased expression of a gene encoding a major siderophore in *Pseudomonas* spp.^41,43^, which, owing to the role of siderophores in iron acquisition, would presumably have significant effects on both competitive ability *and* niche overlap between non-plasmid and plasmid carrying strains.

————–

Together with the preceding simulated scenarios, these examples illustrate the potential insights to be gained via the partitioning of costs of resistance into changes in competitive ability and niche overlap. Thus far in our paper, however, we have ignored the role of temporal variability in mediating competitive ability and the strength of species interactions. From ecological theory ^24,29,31,^ we know that temporal variability can play a significant role in maintaining coexistence when competing species trade-off along a time varying environmental axis (e.g. antibiotic concentration). In addition, we have been deliberately vague in defining competitive ability, beyond the somewhat fuzzy notion of “growing better (or worse)” on available resources (but see Supplementary Information S2 for a mathematically explicit definition). This vagueness obscures a subtle but important property of competitive ability in the context of fitness costs; it depends not only on which resources are available but also on temporal patterns in resource availability, and in particular how they covary through time with antibiotic exposure. In the following section, we turn our attention to the role of constant (or equilibrium) versus fluctuating (non-equilibrium) patterns of antibiotics exposure and resource availability on competitive ability and costs of resistance. Whereas in the preceding the section we considered coexistence at both the intraspecific (i.e. between susceptible and resistant strains) and interspecific level (i.e. between resistant strains and commensal members of the microbiome), we now focus more explicitly on the former.

### Coexistence and resistance in dynamic environments

Microbial communities generally occupy dynamic environments^44^, where fluctuations in environmental factors (e.g. pH, temperature, and of course antimicrobials) and nutrient resources can be expected to drive significant oscillations in population abundances and community composition through time. The pulsed nature of most antibiotic dosing regimes likely acts as a particularly potent driver of variability. Nevertheless, despite a substantial body of literature focusing on the optimisation of dosing regimes (i.e., dose timing and frequency) in the context of pharmacodynamics (e.g.^45^), the implications of dosing regime on the interaction between within-host competition and the evolution of drug resistance have thus far gone largely ignored. Notwithstanding several recent studies that have begun to tackle this question (e.g.^46–49^), as noted by Holmes *et al.*^14^, “the absence of basic knowledge about ideal prescribing regimens represents a significant gap”.

Part of the difficulty in tackling this question is that, in the community context, the efficacy of different dosing regimes should interact closely with the temporal dynamics of limiting nutrients. To our knowledge, there has been little to no research looking explicitly at how the interaction between resource delivery regime (e.g. feeding patterns) and antibiotic exposure regulates the evolution of drug resistance. We can nevertheless draw some insight from emerging research into the effects of feeding patterns, independent of nutritional composition, on gut microbiome stability^50–52^.

Even without exposure to antibiotics, the gut is already a highly dynamic environment^53^, where natural cycles in fasting and feeding generate fluctuations in nutrients, pH and secondary metabolites^50^. A wealth of research has already documented the conspicuous effects diet has on the gut microbiota; in both humans and mice it has already been shown that a change in diet can dramatically shift an individual’s gut microbial community in as little as a few hours^54–56^. Much less attention has been given to understanding how the act and frequency of feeding (e.g., time restricted vs unrestricted), independent of dietary composition or total calorific intake, modulate gut microbiome composition. A small number of recent studies have nevertheless begun to tackle this research question, the findings of which provide compelling evidence that feeding patterns can indeed reshuffle competitive hierarchies within the gut^50–52,57,58^. For example, Zarrinpar et al.^50^ found that mice subjected to the same diet but different feeding patterns differed in their gut microbiota, with natural cycles of feasting and fasting corresponding to predictable changes in microbial composition. Whereas mice allowed to feed ad libitum on a high fat diet exhibited low levels of compositional cycling, mice restricted to night-time feeding on the same diet were characterised by significant cyclical variation in both protective and pathogenic taxa^50^. Thaiss et al.^51^ similarly found intra-daily feeding patterns to be a strong determinant of microbial cycling within the gut, with mice fed exclusively at night or day exhibiting a corresponding phase shift in the peak abundance of particular taxa.

If, as several recent studies have suggested ^7,8,35,59,^ ecological competition can place strong limits on the evolution of antibiotic resistance, it stands to reason that the strength of this inhibitory force will be highly contingent on competitor identity. At the same time, the aforementioned research suggests that subtle differences in feeding pattern alone can have significant impacts on the identity of dominant competitors at time-scales (i.e. hours) that closely correspond to the pharmacological lifespan of antimicrobial drugs. The conventional rationale for prescribing the intake of medicines along with, or in isolation, from food is to either ensure the optimum pharmacological effectiveness of a given drug^60^, or to reduce any potential physiological side-effects^61^. Is it possible that future dosing-dietary interaction guidelines would be best advised to take into account the effect of food consumption on the relative fitness of pathogens and their drug resistant variants in the explicit context of the microbiome? In the following section, we draw on ecological theory to lay out a series of testable predictions for how antibiotic delivery and its interactions with resource availability and resistance trade-offs may be expected to modulate the relative fitness and coexistence of resistant and susceptible strains.

### Predicting competitive outcomes

All else being equal, under constant antibiotic exposure, a resistant strain will exclude a susceptible strain above some critical threshold in antibiotic concentration, and a susceptible strain will exclude a resistant strain below that threshold. The threshold is set by the point at which the fitness costs of resistance and susceptibility cross-over. Most pertinent to the current discussion, however, is that coexistence under this simplistic scenario is not possible, beyond the infinitely unlikely case of neutral coexistence when relative fitness is exactly equal. By contrast, the pulsed delivery (typical of most regimes) of a bacteriostatic antibiotic can promote stable coexistence of resistant and susceptible strains at the same time-averaged concentration that would lead to one phenotype being excluded under constant exposure (Fig 3). This occurs because the pulsing of an antibiotic that impedes growth effectively concentrates intraspecific competition relative to interspecific competition in time; or in the language of coexistence theory, it decreases niche overlap through time via a mechanism community ecologists refer to as a temporal storage effect^62,63^. To emphasize that this represents a stabilizing process (and not the unlikely case of perfectly balanced fitnesses), note that under sufficiently large pulse intervals, a spectrum of fitness costs allow the resistant strain to persist alongside the susceptible strain (grey wedge in Fig 3a).

**Figure 3:**
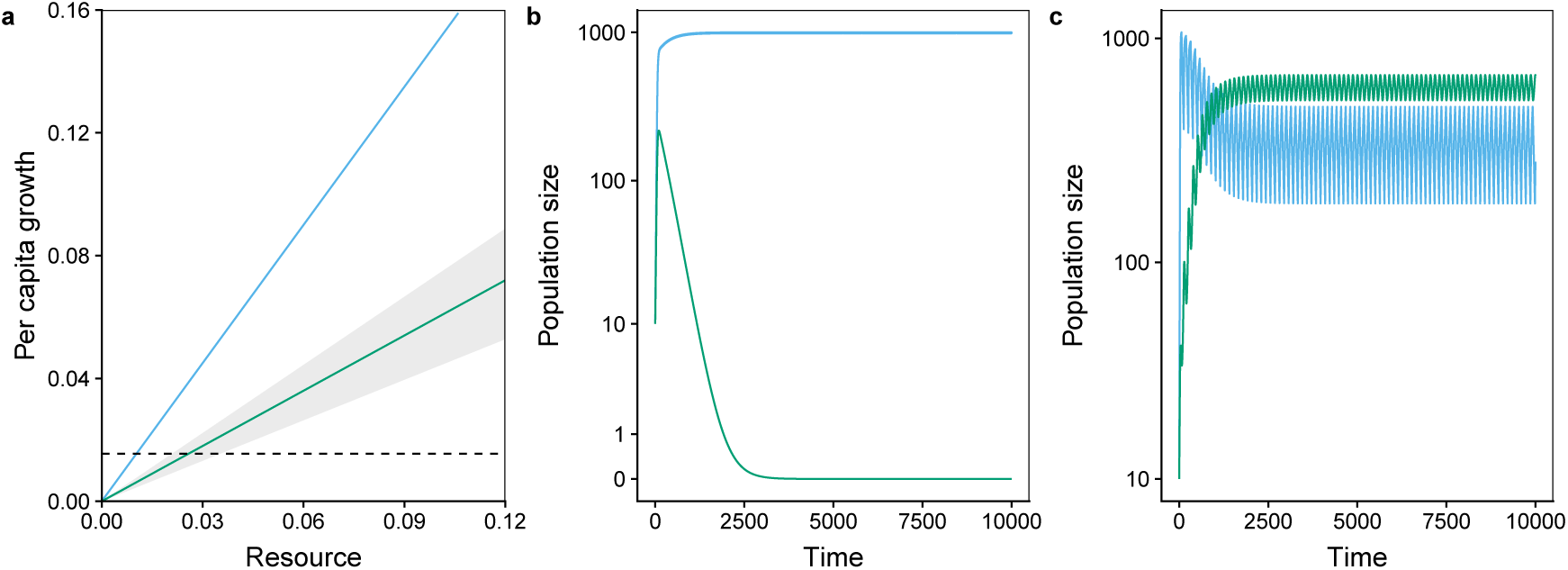
The effect of constant versus pulsed delivery of bacteriostatic antibiotics on coexistence between an susceptible strain (blue) and a resistant mutant (green). **(a)** Per-capita growth responses to a limiting resource under antibiotic-free conditions, with the susceptible strain exhibiting superior growth. Under antibiotic exposure, the susceptible strain is unable to grow (not shown) while the resistant strain is unaffected. **(b)** When antibiotics are delivered at a constant rate below that at which the detrimental effect on growth exceeds the cost of resistance, the susceptible strain excludes the resistant strain. **(c)** Pulsing of antibiotics at the same time-averaged concentration as in **(b)** allows the resistant strain to coexist alongside the susceptible strain. For the given pulse interval, stable coexistence emerges for all costs of resistance spanning the grey shaded area in **(a)**. See Supplementary Information S3 for model simulation design and parameters.

To illustrate this more completely, Figure 4 summarizes simulations of resource competition between a susceptible strain and a resistant strain characterized by varying degrees of fitness costs (decrease in per-capita growth rate) across increasingly large antibiotic pulsing intervals (see Supplementary Information S3 for simulation parameters). The pulsing interval represents switching time between antibiotic-free and antibiotic-exposed conditions. As such, the total (or average) amount of time the competing strains are exposed to an antibiotic is constant across pulsing intervals, but the more frequent the switching, the more constant the bacteria perceive the environment. The main observation we wish to highlight is that when conditions switch rapidly back and forth, only a narrow range of fitness costs allow the resistant strain to persist. However, as the length of the pulse interval increases, the range of fitness costs that enable the resistant strain to persist broadens. In sum, all else being equal, fluctuating conditions generated by the pulsing of antibiotics benefits resistant strains over susceptible strains.

**Figure 4:**
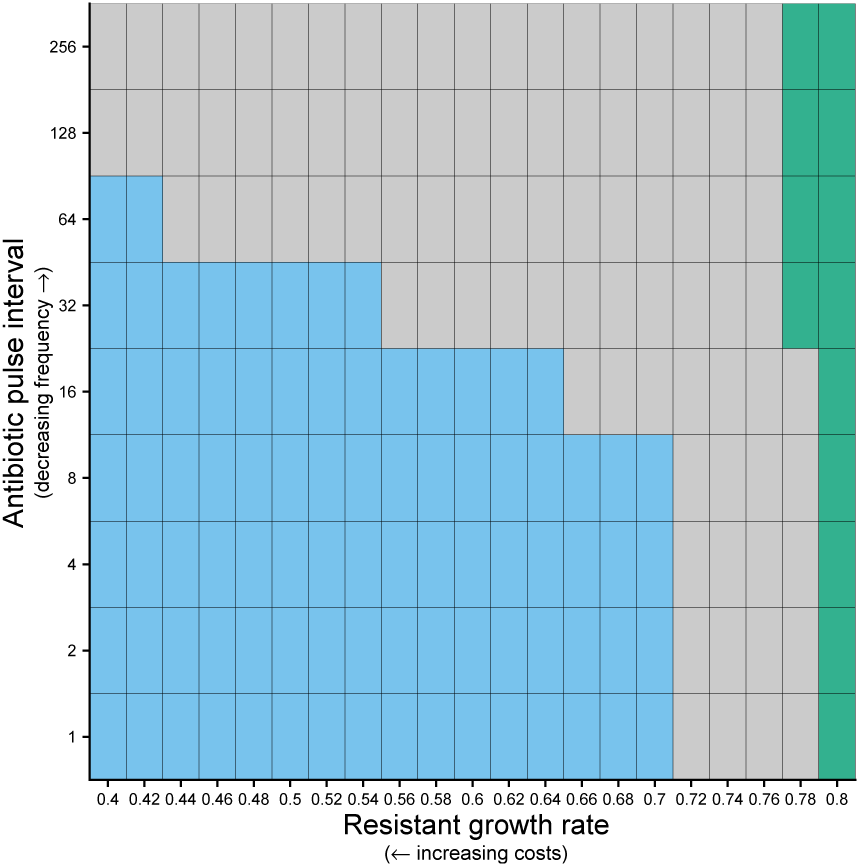
Coexistence and exclusion of a susceptible and resistant strain across antibiotic pulse intervals of increasing length (y-axis) but the same time-averaged concentration; and decreasing costs of resistance (x-axis). In the bottom left corner large costs of resistance coupled with rapid pulsing results in exclusion of the resistant strain (blue cells). In the top right corner small costs of resistance coupled with long pulse intervals results in exclusion of the susceptible strain (green cells). The range of costs of resistance leading to coexistence (grey cells) increases with longer pulse intervals. See Supplementary Information S3 for simulation parameters.

But what if all else isn’t equal? An assumption of the above simulations was that the limiting resource entered the system at a constant rate, as it would in a chemostat. This of course is unlikely in many microbial environments, including, for example, the animal gut where feeding patterns can cause rapid temporal (hourly to daily) fluctuations in the composition of the microbiota^50–53^. Now we wish to relax this assumption to investigate how fluctuations in resource delivery affect coexistence of resistant and susceptible strains.

When resources are delivered constantly, and multiple species are limited by the same resource, the species with the lowest maintenance requirement (R*, resource concentration at which growth rate is zero) will exclude all other competitors^23,25,28^. However, when resources fluctuate through time, it is no longer possible for the species with the lower R* to maintain the resource at a low level and so the competitive advantage of being a resource ‘gleaner’ is dramatically weakened^64^.

Instead, an ‘opportunist’ species with a higher R* but a larger growth rate at high resource levels is more likely to be the superior competitor. Indeed, under the right frequency and amplitude of resource pulses, these two strategies can coexist via a mechanism termed ‘relative nonlinearity of competition’^63–65^. The name of this mechanism refers to the requisite difference in the curvature of species’ growth responses to resource concentration which allows for the opportunist-gleaner trade-off.

The implication of these opposing resource acquisition strategies is that a cost of resistance that is severe in an equilibrium system, may be negligible in a much more variable system and vice versa. Consider an environment in which antibiotics are again pulsed into the system on some periodic schedule (e.g., the oral ingestion of antibiotics in medical treatment or run-off from agricultural application of antibiotics to livestock). Irrespective of where the resistant strain sits along the resource trade-off spectrum, first note that resource pulses that come out of phase with antibiotics should benefit the susceptible strain, while resource pulses that come in phase with the antibiotics will benefit the resistant strain. The effect of pulsing resources anti-phase is that it limits the gains that can be made by the resistant strain when exposure to antibiotics eliminates competition with the susceptible strain, while at the same time giving the susceptible strain the resources it needs when it is the better competitor in the antibiotic free environment. Conversely pulsing resources in phase with antibiotics allows the resistant strain to effectively consume all available resources. But what if resource availability is not correlated with antibiotic pulsing (e.g. if resources are available in equal concentration in the presence and absence of antimicrobials)? In the absence of strong temporal covariance between resources and antibiotics, an interaction between mode of resource delivery (i.e. pulsed vs. constant) and the specific resistance phenotype comes into play.

Consider a drug resistant strain which exhibits the strongest costs of resistance at high concentrations of a limiting resource, i.e., the difference in its growth rate relative to the drug susceptible ancestor is greatest at high resource levels *(Max growth trade-off* in Fig 5). We again make the simple assumption that antibiotics are pulsed into the system on some fixed schedule, such that competing phenotypes experience equal fixed-length periods in antibiotic free and exposed conditions. Now, however, we assume resource pulses are nested (i.e. shorter in length) within antibiotic pulses such that the total amount of resources available is spread equally across the two opposing environmental conditions. In this case we see that at one extreme, small and frequent resource pulses allow the resistant strain to exclude the susceptible strain (green shaded cells in bottom left of main left panel in Fig 5). This is because small frequent pulses minimise the fitness difference between the two phenotypes in the antibiotic free environment. However, with increasingly longer intervals and/or larger resource pulses, the susceptible strain will be able to coexist with the resistant strain (grey shaded cells in the middle of main left panel), until the other extreme is reached and the susceptible strain excludes the resistant strain (blue shaded cells in top right of main left panel). This is because large infrequent pulses maximise the fitness difference between the two phenotypes in the antibiotic free environment. This prediction is in fact consistent with experimental data obtained by Maharjan and Ferenci ^66^ who found that significant fitness differences observed between rifampicin resistant *E. coli* and the ancestral wild type under nutrient rich conditions, largely disappeared when the same strains were competed under nutrient limiting conditions in a chemostat.

**Figure 5:**
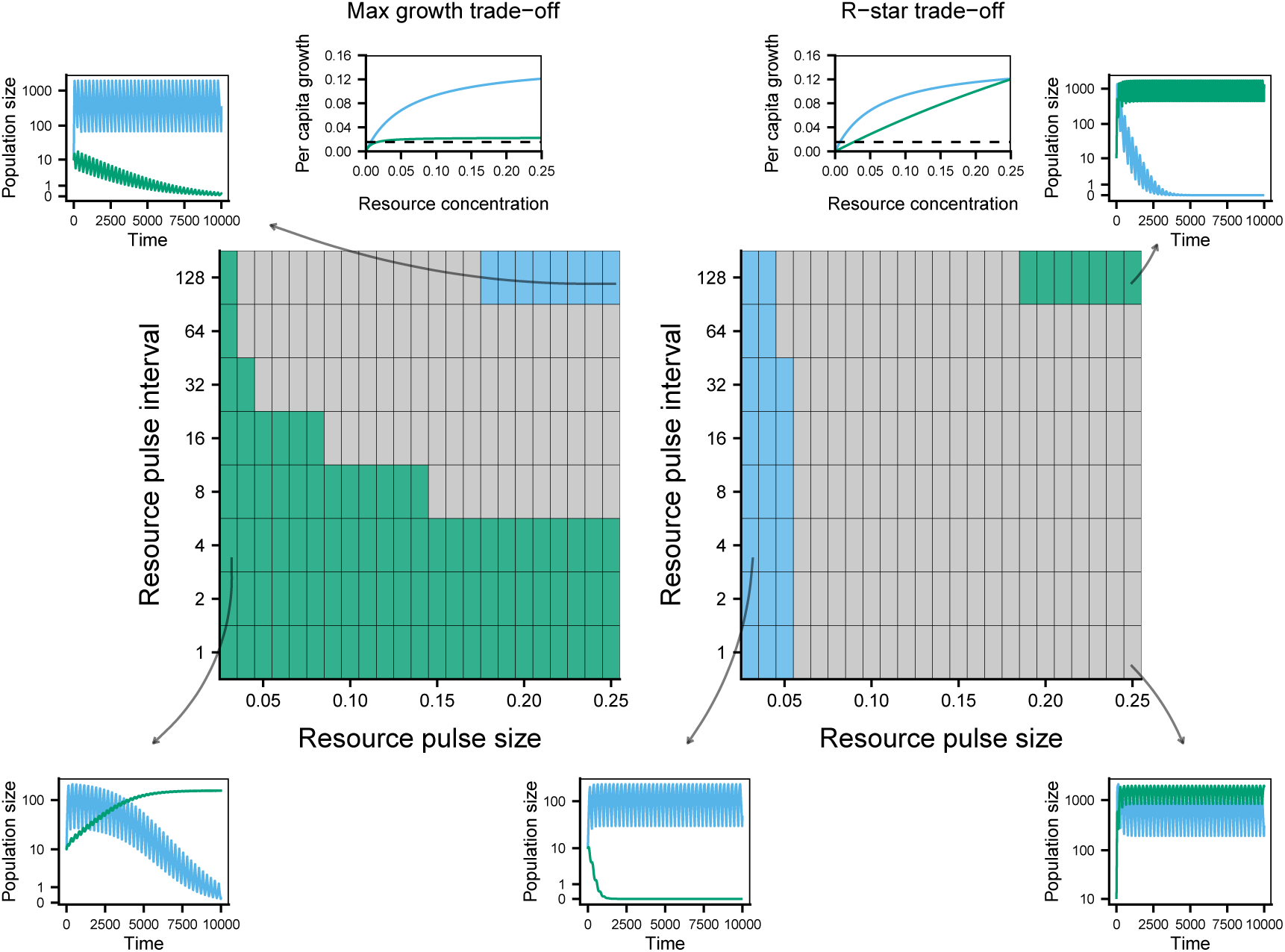
Simulation results illustrating the interactive effects of antibiotic pulsing, resource pulse interval length (y-axis) and resource pulse size (x-axis) on competitive outcomes under different resource-uptake associated costs of resistance (max-growth trade off in left panel; R* trade-off in right panel). The length of each antibiotic-free and antibiotic-exposed interval is fixed at 128 units of time. Total resource availability is equal across the antibiotic-free and antibiotic-exposed intervals. Green cells indicate exclusion of the susceptible strain; blue cells indicate exclusion of the resistant strain; grey cells indicate coexistence. See Supplementary Information S4 for simulation parameters.

Now consider the alternative scenario, where a drug resistant strain exhibits the strongest costs of resistance at low concentrations of a limiting resource, i.e. the difference in its growth rate relative to the drug susceptible ancestor is greatest at low resource levels (*R*^*\**^ *trade-off* in Fig 5).With an *R*^*\**^ trade-off we see the opposite relationship between resource pulse interval/size and community dynamics. In diametric contrast with a max-growth trade-off, small frequent resource pulses allow the susceptible strain to exclude the resistant strain; intermediate pulse intervals/sizes lead to coexistence; while large infrequent pulses, which minimise costs of resistance, help the resistant strain to exclude the susceptible strain.

To reiterate our main message, we see that the implication of a trade-off in competitive ability for resources on costs of resistance can only be understood in light of the governing resource regime. An apparently large cost of resistance observed experimentally in batch culture, which exemplifies pulsed regimes, might disappear under more constant resource regimes. By the same argument, the apparent absence of a large cost of resistance in experimental batch culture could mask significant costs in the absence of resource pulsing.

### Limitations

Perhaps the most conspicuous limitation of much ecological theory in a clinical context, is that the clinical response of interest is typically abundance (i.e. pathogen load) rather than qualitative state (i.e. coexistence vs. exclusion). A great deal of theoretical ecology is grounded in the study of dynamical systems. Classically, the analysis of dynamical systems concentrates on evaluating what system properties (e.g. parameter values or initial values for the state variables) lead to qualitatively different end points. For example, can two competing species coexist at equilibrium, or is one excluded by the other? This is to say, the analysis of dynamical systems is typically less concerned with the relative sizes of the state variables at equilibrium than it is with determining which have non-zero equilibrium values and the stability of those equilibrium values. From the perspective of understanding the effect of ecological interactions on the persistence of drug resistant strains, variation in absolute abundance between stably coexisting strains is of paramount importance; a pathogen strain that persists at low abundance may be of negligible concern to one that persists at high abundance, even though dynamically the systems are qualitatively equivalent. Populations at low abundance also become increasingly vulnerable to demographic stochasticity, and several researchers have argued that the immune system can clear the residual infection once the population is sufficiently small^48^, indicating that the kinds of qualitative indicators derived from dynamical analyses might be less informative in practical terms. Fortunately, efforts are already underway within the theoretical literature to more explicitly incorporate demographic stochasticity into coexistence analysis^67,68^, and there is no constraint on extending the types of models regularly adopted by community ecologists to incorporate the immune response.

The latter concern alludes to a much broader tension between theory and reality - that is the trade-off between generality and complexity. In addition to the immune system, we can make a long list of other ecological and evolutionary factors that may well have an overriding impact on the dynamics identified herein, from predation, cross-feeding, spatial heterogeneity and direct competition (i.e. bacterial warfare), to compensatory mutations and horizontal gene transfer. Again there is no limitation to the incorporation of these complexities into the kinds of models we have used for our analyses, and we certainly encourage studying dynamics under these more complex scenarios (see Future Directions). Nevertheless, from an ecological perspective, resource competition arguably is the most pervasive and canonical interaction type, at least within trophic levels. Furthermore, owing to energy limitations, anaerobic habitats such as the mammalian gut are thought to support comparatively flat ecosystems, limited to at most two trophic levels with very few secondary consumers ^44^. Combined with the apparent sparsity of research into the ecology of antibiotic-resource interactions, we believe this provides a strong rationale for renewed attention on the role of resource competition in mediating the evolution of antibiotic resistance.

### Future Directions

Experimental tests of the ideas presented represent a natural jumping-off point for future research. The nature and scope of these experiments can be usefully separated into those which explore the partitioning of costs of resistance into niche overlap and competitive ability differences, and those that explore the relative timing of antibiotic and resource pulses on the persistence of antibiotic resistance.

A central question deriving from these analyses is to what extent costs of resistance are disproportionately captured by changes in niche overlap versus changes in competitive ability, and to what extent this balance is contingent on the focal taxa, the type of antibiotic, and/or the nature of the resistance mutation. The most convenient approach to the partition is to empirically parameterise a phenomenological competition model (e.g. Lotka-Volterra), from which niche overlap and competitive ability differences can be quantified (see Box 1)^28,69^. For high resolution time series data, the Lotka-Volterra model can be parameterised via statistical fits of the dynamical model (a system of ordinary differential equations)^70^. An alternative approach, which is common in the plant ecology literature, is to obtain the competition coefficients directly based on statistical fits of per-capita growth rate at varying densities of the focal and non-focal strain^71–73^. The latter approach, however, will only be accurate over a short enough time-interval such that the independent variable, population density, is not allowed sufficient time to grow or decay by a large amount.

As recognised by Holmes *et al.*^14^, it remains surprisingly unclear how different prescribing regimes affect the evolution of antibiotic resistance. Testing the relevant theoretical predictions presented in the current work would require an experimental setup in which the delivery of antibiotics and resources can be regulated through time from being continuous at one end of the spectrum to highly pulsed at the other. For *in vitro* work this almost certainly necessitates a chemostat setup, although a semi-continuous serial transfer approach may be be sufficient under high transfer rates as facilitated by liquid handling robotics. Looking further forward, there are also opportunities to test these ideas *in vivo* using model systems such as mice or *Drosophila* where antibiotic delivery and food availability can be tightly regulated.

The experimental approaches outlined in the preceding paragraph all undoubtedly include significant infrastructural overheads, which may explain the paucity of experimental research into the effect of different prescribing regimes on the evolution of resistance. An alternative approach with a lower infrastructural barrier to entry would be to start by simply quantifying the functional nature of resource trade-offs associated with costs of resistance. Which is more common, a loss of competitive ability at high or low resource levels (see e.g.^66^)? And does this depend on the limiting resource, the focal taxa, the antibiotic, or the resistance mutation? This information would enable investigators to parameterise more mechanistic models of resource-competition to explore and make predictions on the effects of different prescribing regimes on resistance evolution.

Beyond empirical work, as already indicated, there is substantial scope for theoretical and computational studies that incorporate additional ecological and evolutionary complexities into these analyses. With respect to ecological theory, there already exists a substantial body of theory aimed at incorporating the effects of predation into the partition of niche overlap and competitive ability^69^. For example, apparent competition, where the effect of a shared predator mimics the role of resource competition, can be mathematically partitioned into the separate coexistence mediating components. This could be particularly relevant to understanding the combined effects of resource competition and parasitism by bacteriophages. Infection of pathogenic bacteria by bacteriophages has been indicated as a significant factor regulating antibiotic resistance owing to putative trade-offs in resistance to antibiotics and phages^74^. Few theoretical studies have considered the effect on coexistence of other ecological processes that are thought to be a common feature of microbial systems, such as cross-feeding and direct competition but see ^75^, but it is again straightforward to incorporate these dynamics into classic models of resource competition. Similarly, the theoretical literature on antibiotic resistance should provide a wealth of examples for how other evolutionary and physiological factors, including horizontal gene transfer, mutation rates and an active immune system, might be incorporated into the kinds of ecological models we have focused on here.

## Conclusions

The motivation behind this article was to present recent concepts from theoretical ecology that could prove useful in understanding the evolution of antibiotic resistance and the coexistence of susceptible and resistant pathogens in microbial communities. It remains to be seen whether these insights can ultimately be of use in developing strategies to manage resistance. As with much theoretical knowledge, the path to generality is littered with simplifying assumptions, which still need to be tested and verified empirically. Nevertheless, with increasing awareness that drug resistant pathogens are embedded within complex species interaction networks, it would be short-sighted not to take advantage of a wealth of valuable theory and models at the frontier of community ecology.

## Acknowledgements

The authors thank David McLeod, Sebastian Bonhoeffer and the Pathogen Ecology group at ETH for thoughtful discussions and comments on an earlier draft of this manuscript.

## Supporting Information

Additional supporting information may be found in the online version of this article:

## Footnotes

† Differences in competitive ability are also commonly referred to as fitness differences, but we adopt the former terminology here to avoid confusion with evolutionary fitness, which is the combined outcome of differences in competitive ability and niche overlap.

## Notes

### Competing Interest Statement

The authors have declared no competing interest.

