## Supplementary Information S1-S4 for "Antibiotics in microbial communities: an ecological frame of resistance"

### Supplementary Information:
### Antibiotics in microbial communities: an ecological frame of resistance


Andrew D. Letten*[1,2], Alex Hall[2], and Jonathan Levine[3]

[1]School of Biological Sciences, University of Queensland, Brisbane, Queensland 4072, Australia
[2]Institute of Integrative Biology, Department of Environmental Systems Science, ETH Zürich, 8092 Zürich, Switzerland
[3]Princeton University, USA


May 11, 2020

**Supplementary information S1: Obtaining mechanistic definitions for competitive ability and niche differences**

In order to obtain mechanistic definitions of competitive ability and niche differences, we start with a classic mechanistic model of competition for two consumers feeding on two resources,

$$\frac{\mathrm{d}N_i}{\mathrm{d}t} = N_i(\mu_{ia}R_a + \mu_{ib}R_b - m_i), \tag{S1}$$

$$\frac{\mathrm{d}N_j}{\mathrm{d}t} = N_i(\mu_{ja}R_a + \mu_{jb}R_b - m_j), \tag{S2}$$

$$\frac{\mathrm{d}R_a}{\mathrm{d}t} = r_a R_a(a - \frac{R_a}{K_a}) - \mu_{ia}R_a q_{ia} N_i - \mu_{ja} R_a q_{ja} N_j, \tag{S3}$$

$$\frac{\mathrm{d}R_b}{\mathrm{d}t} = r_b R_b(a - \frac{R_b}{K_b}) - \mu_{ib}R_b q_{ib} N_i - \mu_{jb} R_b q_{jb} N_j, \tag{S4}$$

where $N_i$ is the population density of consumer $i$, $R_a$ is the density/concentration of resource $a$, $\mu_{ia}$ is the per capita uptake rate of consumer $i$ on $R_a$, $m_i$ is the per capita mortality rate of consumer $i$, $q_{ia}$ is the resource quota (amount of resource per individual) for consumer $i$ on resource $a$, $r_a$ is the intrinsic rate of increase of resource $a$ and $K_a$ is the carrying capacity of resource $a$.

If we make the assumption that the resource dynamics are much faster than the consumer dynamics, we can solve for the equilibrium vales of $R_a$ and $R_b$ by setting Eqs 3 & 4 = 0 and substitute the solutions back into Eqs 1 & 2. We then rearrange the reduced system of equations into a Lotka-Volttera form in order to obtain the mechanistic derivations of the Lotka-Volterra parameters,

$$\begin{aligned}\frac{\mathrm{d}N_i}{\mathrm{d}t} = N_i((&\mu_{ia}K_a + \mu_{ib}K_b - m_i) - \\ &(\frac{\mu_{ia}^2 q_{ia} K_a}{r_a} + \frac{\mu_{ib}^2 q_{ib} K_b}{r_b})N_i - \\ &(\frac{\mu_{ia}\mu_{ja} q_{ja} K_a}{r_a} + \frac{\mu_{ib}\mu_{jb} q_{jb} K_b}{r_b})N_j),\end{aligned} \tag{S5}$$

$$\begin{aligned}\frac{\mathrm{d}N_j}{\mathrm{d}t} = N_j((&\mu_{jb}K_b + \mu_{ja}K_a - m_j) - \\ &(\frac{\mu_{jb}^2 q_{jb} K_b}{r_b} + \frac{\mu_{ja}^2 q_{ja} K_a}{r_a})N_j - \\ &(\frac{\mu_{jb}\mu_{ib} q_{ib} K_b}{r_b} + \frac{\mu_{ja}\mu_{ia} q_{ia} K_a}{r_a})N_i).\end{aligned} \tag{S6}$$

The Lotka-Volterra parameters can then in turn can be substituted into the formulas for competitive ability differences and niche overlap given in Box 1 in the main text, such that:

$$\rho = \sqrt{\frac{(\frac{\mu_{ia}\mu_{ja}q_{ja}K_a}{r_a} + \frac{\mu_{ib}\mu_{jb}q_{jb}K_b}{r_b}).(\frac{\mu_{jb}\mu_{ib}q_{ib}K_b}{r_b} + \frac{\mu_{ja}\mu_{ia}q_{ia}K_a}{r_a})}{(\frac{\mu_{ia}^2 q_{ia}K_a}{r_a} + \frac{\mu_{ib}^2 q_{ib}K_b}{r_b}).(\frac{\mu_{jb}^2 q_{jb}K_b}{r_b} + \frac{\mu_{ja}^2 q_{ja}K_a}{r_a})}}, \tag{S7}$$

$$\frac{k_j}{k_i} = \frac{\mu_{jb}K_b + \mu_{ja}K_a - m_j}{\mu_{ia}K_a + \mu_{ib}K_b - m_i}\sqrt{\frac{(\frac{\mu_{ia}^2 q_{ia}K_a}{r_a} + \frac{\mu_{ib}^2 q_{ib}K_b}{r_b}).(\frac{\mu_{ia}\mu_{ja}q_{ja}K_a}{r_a} + \frac{\mu_{ib}\mu_{jb}q_{jb}K_b}{r_b})}{(\frac{\mu_{jb}^2 q_{jb}K_b}{r_b} + \frac{\mu_{ja}^2 q_{ja}K_a}{r_a}).(\frac{\mu_{jb}\mu_{ib}q_{ib}K_b}{r_b} + \frac{\mu_{ja}\mu_{ia}q_{ia}K_a}{r_a})}}. \tag{S8}$$

**Supplementary information S2: Model parameters: Figures 1 & 2**

*Resource parameters*
$K_a = K_b = 1$, $r_a = r_b = 1$.

*Growth parameters*
Sensitive strain: $\mu_{sens,a} = 0.283, \mu_{sens,b} = 0.283, m_{sens} = 0.0828, q_{sens,a} = q_{sens,b} = 0.01$
Resistant strain (i): $\mu_{res_i,a} = 0.2, \mu_{res_i,b} = 0.2, m_{res_i} = 0.2, q_{res_i,a} = q_{res_i,b} = 0.01$
Resistant strain (ii): $\mu_{res_{ii},a} = 0, \mu_{res_{ii},b} = 0.283, m_{res_{ii}} = 0.0828, q_{res_{ii},a} = q_{res_{ii},b} = 0.01$
Commensal strain: $\mu_{com,a} = 0, \mu_{com,b} = 0.8, m_{com} = 0.0828, q_{com,a} = q_{com,b} = 0.01$

Note that mortality rates are kept the same ($m = 0.0828$) for all strains except resistant $i$ ($m = 0.2$). Accompanying a decrease in growth on both resources with an increase in mortality in resistant $i$ is necessary to ensure the two resistant phenotypes have equal intrinsic rates of increase and carrying capacities (assuming no change in niche overlap). In the absence of a change in mortality (or niche overlap), the two resistant phenotypes can still have the same integrated competitive ability, but will have different combinations for their logistic growth parameters ($r$ & $\alpha_{ii}$).

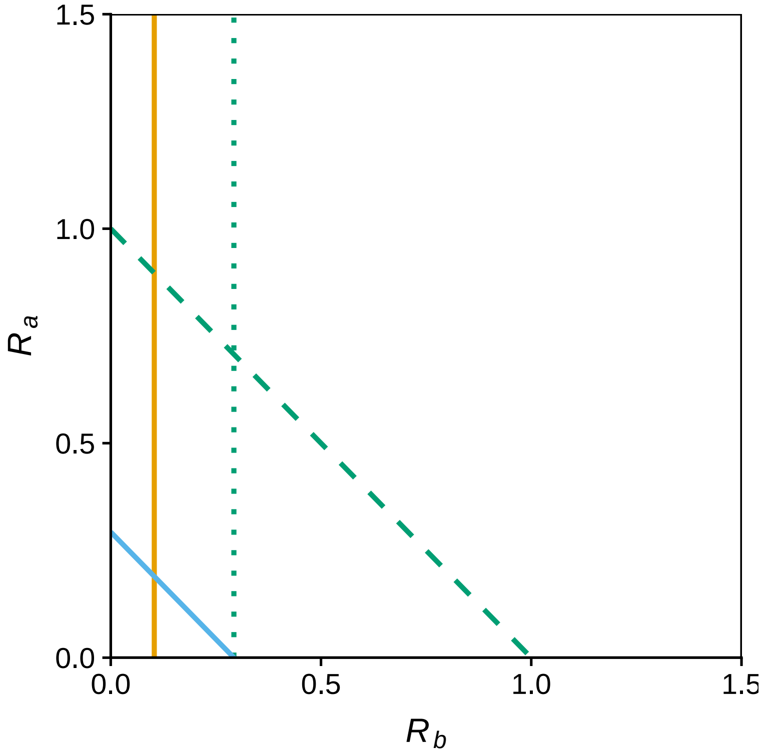


**Figure S1:** Strain resource use profiles in the resource phase plane. Solid, dashed and dotted lines denote the zero net growth isoclines for each strain: sensitive (blue), resistant i (dashed green), resistant ii (dotted green), commensal (orange).

#### Supplementary information S3: Simulating competition under antibiotic fluctuations

In order to explore the effect of antimicrobial pulsing on the relative fitness of resistant and sensitive strains, we modified the above consumer-resource model to include two consumers (sensitive and resistant) competing for a single resource under continuous supply (i.e. chemostat dynamic).

$$\frac{\mathrm{d}N_r}{\mathrm{d}t} = N_r(\mu_r R - m) \tag{S9}$$

$$\frac{\mathrm{d}N_s}{\mathrm{d}t} = N_s(\mu_s(A)R - m) \tag{S10}$$

$$\frac{\mathrm{d}R}{\mathrm{d}t} = d(S - R) - \mu_r R q_r N_r - \mu_s(A) R q_s N_s \tag{S11}$$

where $d$ is the resource inflow/outflow rate, $S$ is the resource supply concentration, and $\mu_s(A)$ (resource uptake rate of the sensitive strain) is a constant in the presence of the antibiotic and 0 in its absence.

**Model parameters: Figure 3**

In Figure 3, the interval between antibiotic exposed and antibiotic free conditions was fixed at 64 timesteps each.

*Resource parameters*
$S = 0.12, d = 0.01552$

*Growth parameters*
$m = 0.01552$
Sensitive strain (antibiotic free conditions): $\mu_s = 1.5, q_s = 0.0001$
Resistant strain: $\mu_r = 0.6, q_r = 0.0001$

**Model parameters: Figure 4**

In Figure 4, model parameters were varied factorially across nine pulse interval lengths (1, 2, 4, 8, 16, 32, 64, 128, 256 timesteps) and 21 costs of resistance (min $\mu_r = 0.4$, max $\mu_r = 0.8$, increment = 0.02). All other parameters as above.

**Supplementary information S4: Simulating competition under antibiotic and resource fluctuations**

In order to explore the effect of variation in the timing of antimicrobial and resource pulsing on the relative fitness of resistant and sensitive strains, we modified S9-S11 such that the consumer functional responses are given by the Monod equation, and the resources are delivered in discrete pulses:

$$\frac{\mathrm{d}N_r}{\mathrm{d}t} = N_r\left(\frac{\mu_{max_r} R}{k_r + R} - m\right) \tag{S12}$$

$$\frac{\mathrm{d}N_s}{\mathrm{d}t} = N_s\left(\frac{\mu_{max_s}(A) R}{k_s + R} - m\right) \tag{S13}$$

$$\frac{\mathrm{d}R}{\mathrm{d}t} = -dR - \frac{\mu_{max_r} R}{k_r + R} q_r N_r - \frac{\mu_{max_s}(A) R}{k_s + R} q_s N_s, t \neq k\tau, k = 1, 2, ..., \tag{S14}$$

$$\Delta R = R_0 + R, t = k\tau, k = 1, 2, ..., \tag{S15}$$

where $\mu_{max}$ is the maximum growth rate of the consumer, $k$ is the half saturation constant (resource concentration at which growth is half $\mu_{max}$), $R_0$ is the size of the resource pulse, and $\tau$ is the time interval between resource pulses. $\mu_{max_s}(A)$ (maximum growth rate of the sensitive strain) is a constant in the presence of the antibiotic and 0 in its absence.

##### Model parameters: Figure 5

In Figure 5, the interval between antibiotic exposed and antibiotic free conditions was fixed at 128 timesteps, and the resource parameters were varied factorially across eight resource pulse frequencies ($\tau = 1, 2, 4, 8, 16, 32, 64, 128$ timesteps) and 23 resource pulse sizes (min-$\Delta R = 0.03$, max-$\Delta R = 0.25$, increment = 0.01).

*Growth parameters*

Sensitive strain (antibiotic free conditions): $\mu_s = 1.5, q_s = 0.01, q_s = 0.0001$
Resistant strain: $\mu_r = 0.6, q_r = 0.0001$
